# Transcriptomic forecasting with neural ODEs

**DOI:** 10.1101/2022.08.04.502825

**Authors:** Rossin Erbe, Genevieve Stein-O’Brien, Elana J. Fertig

## Abstract

Single cell transcriptomics technologies can uncover changes in the molecular states that underlie cellular phenotypes. However, understanding the dynamic cellular processes requires extending from inferring trajectories from snapshots of cellular states to estimating temporal changes in cellular gene expression. To address this challenge, we have developed a neural ordinary differential equation-based method, RNAForecaster, for predicting gene expression states in single cells for multiple future time steps in an embedding-independent manner. We demonstrate that RNAForecaster can accurately predict future expression states in simulated single cell transcriptomic data with cellular tracking over time. We then show that using metabolic labeling scRNA-seq data from constitutively dividing cells, RNAForecaster accurately recapitulates many of the expected changes in gene expression during progression through the cell cycle over a three day period. Thus, RNAForecaster enables short term estimation of future expression states in biological systems from high-throughput datasets with temporal information.

## Introduction

Cells are dynamic and constantly changing. Predicting their future molecular states enables greater understanding of how biological systems will change naturally and in response to perturbation. A limitation of single cell RNA sequencing (scRNA-seq) technologies is that they destroy the cell to measure its molecular state. Therefore, scRNA-seq cannot track the specific molecular trajectory of an individual cell over time. Rather, scRNA-seq yields statistical samples from populations of cells. Performing additional time course experiments can increase the information available about cellular dynamics and cell state changes over time in a biological process. Time course designs can provide substantial information about dynamics of a biological system of interest, but are costly and limited to the time period over which they are measured. While many single-cell technologies do not dynamically profile the molecular state of an individual cell, new metabolic labeling technologies and live cell imaging methods are emerging that are starting to unlock the potential for longitudinal sampling of the molecular states of cells. As these technologies develop, new computational algorithms are needed to determine the distinct transcriptomic states each cell occupied in the past and estimate how cell states will evolve.

Predicting cellular dynamics requires models of both cellular phenotypes and their underlying molecular states. Trajectory inference methods have been widely applied to scRNA-seq data to estimate transitions between cell states (Saelens et al., 2019). Building on the foundation of pseudotime, trajectory inference methods infer the ordering of cells based upon the relative distance of their expression profiles, often incorporating information from low dimensional embeddings or distance between subgroups to define a trajectory of cellular dynamics (Trapnell et al., 2014), (Reid and Wernisch, 2016), (Saelens et al., 2019). These algorithms have been applied to scRNA-seq data collected at different time points (Trapnell et al., 2014), (Reid and Wernisch, 2016), (Schiebinger et al., 2019), along a developmental trajectory (Chen et al., 2019), and through disease states (Campbell and Yau, 2018). Another form of trajectory inference uses optimal transport methods to order cells along a time course by calculating the shortest path in expression space between cell states in a Waddington landscape (Schiebinger et al., 2019), optimal transport with neural ordinary differential equations (Tong et al., 2020), or optimal transport modeled using Jordan-Kinderlehrer-Otto flow learned by an input convex neural network (Bunne et al., 2022). Notably, trajectory inference methods are focused solely on ordering cells and require further extensions to model the gene expression values of these cells forward or backward in time or to account for tracing of individual cells.

RNA velocity, rather than focusing on ordering cells, investigates dynamic cellular processes by estimating the change in expression occurring in each cell based upon the ratio of spliced to unspliced transcripts (La Manno et al., 2018), (Bergen et al., 2020). These methods are most commonly applied to overlay predicted steady state of gene expression onto embeddings, and thus model changes in cellular phenotypes. Velocity methods have been extended to additionally estimate the cellular direction at the level of translation, by comparing spliced counts with protein data, called protein acceleration (Gorin et al., 2020). However, these RNA velocity and protein acceleration methods do not make predictions about future or past cell states beyond the immediate changes in expression. To predict expression state changes further into the future, vector fields have been applied to the concept of RNA velocity to allow for prediction of future states (Qiu et al., 2022), (Chen et al., 2022). One of these methods, called Dynamo, additionally suggests the use of metabolic labeling scRNAseq variants (Battich et al., 2020), (Qiu et al., 2020), (Hendriks et al., 2019), (Erhard et al., 2019), (Cao et al., 2020), in which cells are treated with a modified uridine for a set period of time before they are harvested for sequencing. This modified uridine is incorporated into the RNAs produced in that labeling period, which allows more recently produced transcripts to be distinguished from older ones. While RNA velocity vector fields methods predict the future cellular expression states of the cells in the data for multiple time steps into the future, they requires that all predicted future states fall within the UMAP or gene-dimensional expression space observed in the data set (Qiu et al., 2022), meaning unseen cell expression states cannot be identified.

To estimate the future transcriptomic states of single cells with dynamic measurements, we have developed a neural ordinary differential equation (neural ODE) (Chen et al., 2018) based method, RNAForecaster. RNAForecaster uses count data as input from two time points in the same cell. This sort of data is not available when using standard scRNAseq protocols, but can be provided using labeled and unlabeled counts from metabolic labeling transcriptomic profiling techniques such as scEU-seq (Battich et al., 2020). The counts from the earlier point in time are provided to the input layer of the neural network, which attempts to predict the expression of each gene at the later time point. This prediction is compared to the actual expression at the later time point to train the network. In metabolic labeling data, where the length of the labeling period is known, this allows for the network to forecast expression in real time. The key distinguishing feature of RNAForecaster from trajectory and RNA velocity methods is that it does not depend on a particular lower dimensional embedding of the data but takes input in the gene dimensional space. Therefore, RNAForecaster does not limit its predictions of future transcriptomic states to the expression space of the input data and attempts to generalize beyond the expression values observed in training. Specifically, training the method on expression values for each gene instead of relying on an embedding of the data provides the potential for this method to predict previously unseen transcriptomic states over a limited time period.

We demonstrate the predictive accuracy of RNAForecaster in simulated data, where we can establish a ground truth regarding future cellular expression states. We then apply RNAForecaster to scEU-seq from constitutively dividing cells and demonstrate that the model can predict the transcriptomic direction of cell cycle progression each hour for three days after the initial expression state is provided to RNAForecaster. Altogether, these analyses demonstrate the utility of neural ODEs for short term forecasting of future expression states from temporally resolved single-cell data.

## Results

### RNAForecaster is a neural ODE based method for predicting future transcriptomic states

We designed RNAForecaster as a neural ODE (Chen et al., 2018) method that can leverage single cell transcriptomics profiling methods with temporal resolution to learn to predict future time points over limited time periods. To enable this analysis, the input to RNAForecaster is two single cell RNA count matrices. The model does not depend on the source of these matrices, only that they measure the same genes and cells in two adjacent time points (Figure 1A). RNAForecaster requires that these two matrices measure the same cell in each column, rather than similar cells harvested separately for sequencing. This is required because RNAForecaster attempts to estimate the future transcriptomic states of each individual cell. We denote the first gene expression matrix as time point t = 0 and the second as time point t = 1. These matrices are used to train a neural ODE.

**Figure 1.**
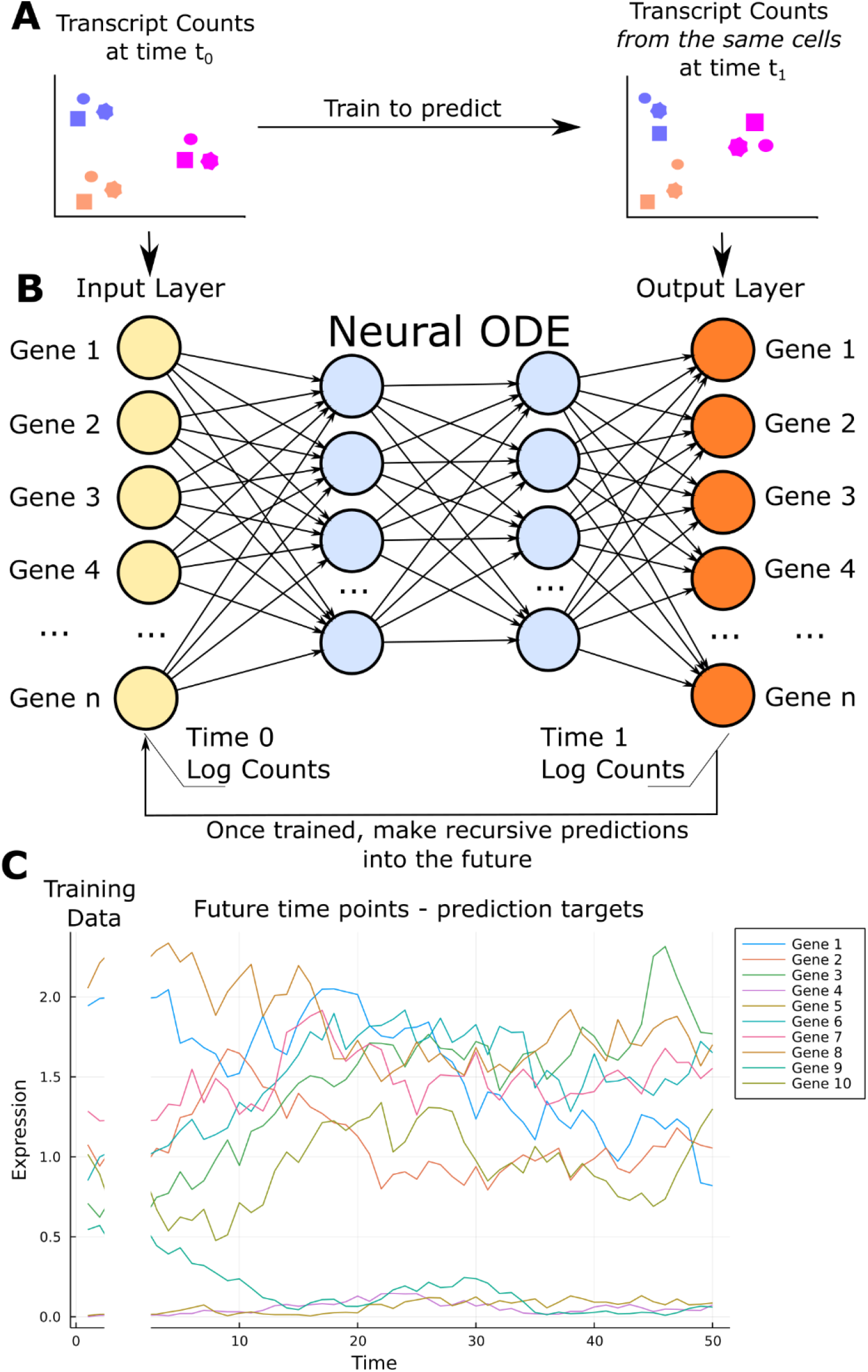
Diagram of RNAForecaster **A** Two count matrices are input to RNAForecaster, each containing the same genes and cells. The counts matrices are from adjacent time points from the same cells, labeled here as t=0 and t=1. **B** The t=0 counts for each cell are input to the input layer of a neural network. The output layer of the neural network has the same number of nodes as the input layer and is compared to the results from the same cell at t=1. The mean squared error between the two forms the loss function which is trained on using an ODE solver to produce a neural ODE. Once the network is trained the output can be fed into the input layer, allowing for prediction of the expression levels at the next time point, which can be repeated recursively to predict for t time steps. **C** A simulation of the expression levels in a cell, showing ten genes over fifty time points. RNAForecaster is trained on the first two time points, using multiple cells in order to learn some generalization of the temporal dynamics between genes. RNAForecaster then attempts to estimate expression of each gene at the later time points.

The training process begins with each cell from the matrix of data from time t = 0 forming an input vector, where the log counts for each gene fill one node in the input layer (Figure 1B). The weights connecting the nodes of the hidden layer(s) and the output layer create an activation function. The output of the activation function represents a prediction of each gene’s expression at time point t = 1. These predictions are then compared with the actual expression level of each gene at time t = 1 based on that input matrix. The mean squared error (MSE) between these values is the loss function of the network. As opposed to standard neural network implementations, weights are updated differently in a neural ODE. Specifically, backpropagation is performed using an ordinary differential equations solver, allowing the network to have a continuous depth and constant memory cost. Thus, the network can yield performant predictions without using many hidden layers and maintains a constant memory requirement, making it computationally cheaper to train than most deep learning alternatives (Chen et al., 2018). After the network is trained, the predicted expression values from the output layer can be fed back into the input layer (Figure 1B), allowing the network to predict the cellular transcriptional state at future time steps. These predictions can be repeated recursively until an arbitrary time t = n, although the propagation of error with each step will cause the prediction error to generally increase over time.

The use of the ODE solver for backpropagation explicitly models dynamical systems, such as the evolution of gene expression values over time, making neural ODEs particularly well suited to predicting future transcriptional states. Additionally, as the neural network does not require a large number of layers to be performant (Chen et al., 2018) it is able to solve this prediction task in a computationally tractable manner. This is a critical feature because using thousands of variable genes as input creates a very large number of network parameters, which would produce a very computationally demanding network to train using other deep neural network architectures. Further, neural ODEs have been found to be particularly accurate at time series predictions relative to other neural network variations (Chen et al., 2018).

To illustrate the prediction task RNAForecaster performs, we provide an example of a sample cell with ten genes (Figure 1C). RNAForecaster is trained on the first two time points from this cell. By training RNAForecaster on many similar simulated cells, each with two time points from the same cell, it can learn the relationships between genes and generalize to make future predictions beyond the gene expression space it was trained on. This challenge of generalizing to a diverse array of transcriptomic states and determination of the temporal limits of predictability will be the focus of the applications we discuss.

### RNAForecaster makes accurate predictions in future expression data outside its training set in simulated single cell transcriptomic data

We generate simulated data to benchmark the feasibility of estimating future transcript counts with RNAForecaster. Simulated temporally resolved single cell expression data was generated using BoolODE (Pratapa et al., 2020) as described in the methods. Briefly, this algorithm simulates gene expression from a system of ordinary differential equations from a known gene regulatory network and incorporates a model that allows for transcriptional busting, and thus contains stochastic elements. To recapitulate the way in which expression data provides a single time snapshot of gene expression, BoolODE simulates a cell’s expression at hundreds of time points and then samples one for inclusion in the output counts matrix. Here, we leverage these additional future expression states as the ground truth for comparison with the predictions of RNAForecaster. To create each simulated data set, we generated a random ten gene network of regulatory relationships between genes. We generated over one hundred randomly generated networks, each of which was used to simulate a single-cell data set with 2000 cells and ten genes at 801 simulated time points.

We begin by training RNAForecaster on two simulated time points, each from the same cell. Standard scRNA-seq data sets are not able to produce this type of training data, but these simulated data allow for a validation of the general principle underlying RNAForecaster. A set of cells is randomly selected as the training set (80%) and the remaining cells form a validation set. We additionally trained a five hidden layer multilayer perceptron (MLP) for comparison on the same data. This MLP model is a feed-forward neural network that can provide a comparison for prediction accuracy using a simple network architecture (see methods for details). The MLP is used as a comparison to benchmark the performance of the neural ODE against the most standard neural network architecture. We first compare predictions of expression at t = 1 in the held out validation set using the expression at t = 0 as input. The RNAForecaster neural ODE significantly outperforms the MLP model on the validation data (p < 1e-16) (Figure 2A), though both methods accurately predict the first time point, with the average mean squared error (MSE) across simulations below 0.015 for both networks.

**Figure 2.**
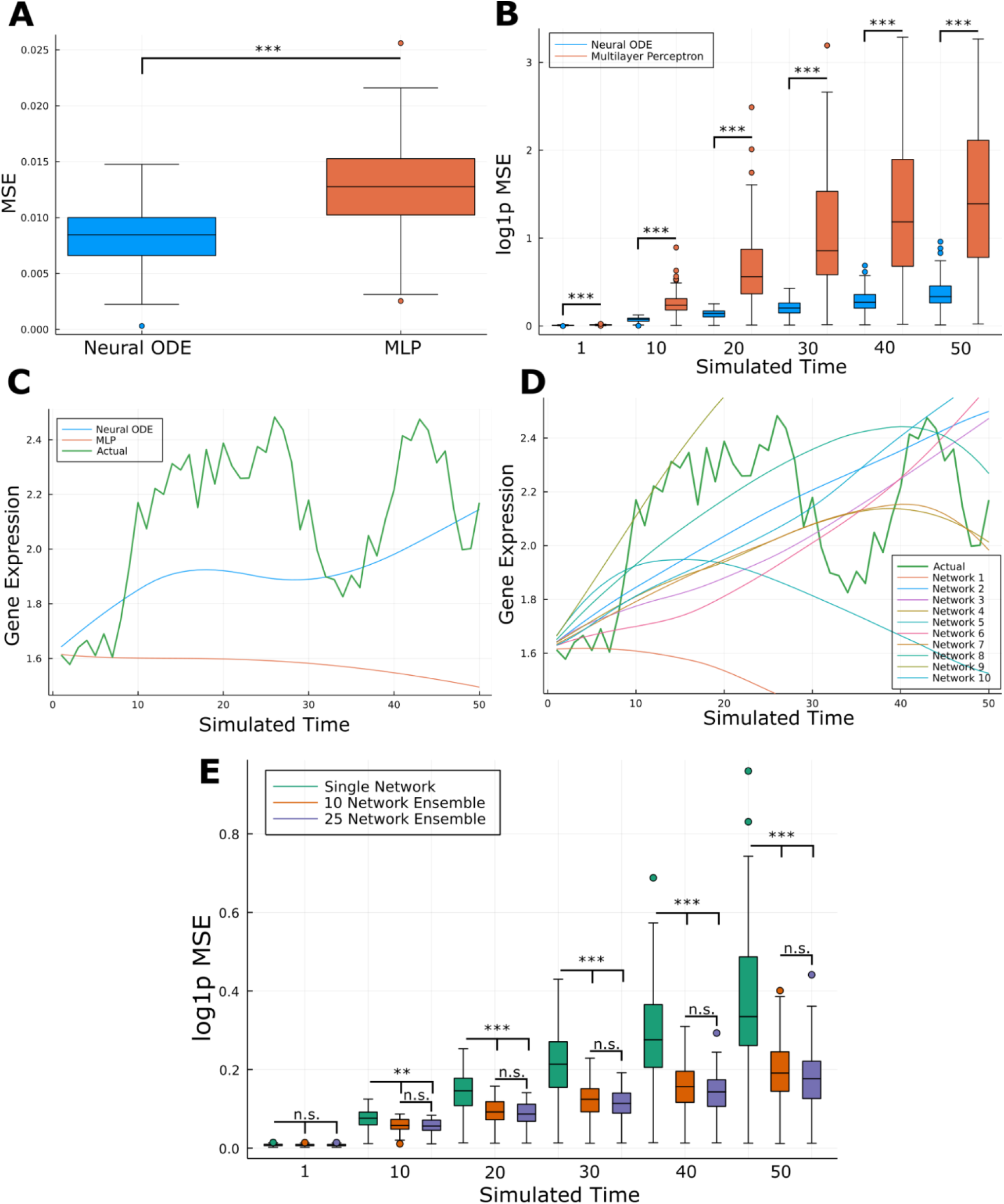
RNAForecaster prediction accuracy in simulated single cell expression data **A** Comparison of MSE loss on the 20% held out validation set of predictions from t=0 to t=1 between a neural ODE and a 5 hidden layer MLP, over all simulations. **B** Comparison of log MSE loss on the next 50 simulated time points between a neural ODE and a 5 hidden layer MLP. **C** A median example of expression prediction of a single gene in a single cell. The predictions of the neuralODE and MLP are shown. **D** The predictions of ten different neural ODEs, each trained using a different initialization of stochastic gradient descent, for the same gene and cell as C. **E** Log MSE loss comparison between a single network neural ODE vs the median predictions from a 10 or 25 network ensemble of neural ODEs. ** p < 1e-6 *** p<1e-10

To determine the temporal range over which predictions can be made, we then tested the ability of the models to predict simulated expression for the next fifty time points for each cell. While the error in both methods increases over time, the neural ODE outperforms the MLP model significantly at all time points (p < 1e-11) (Figure 2B). Error propagates more quickly in the MLP and we additionally observe the presence of extreme outliers in the MLP predictions as early as t = 10, that fall within the same range of the worst predictions of the neural ODE at t = 50. The presence of these inaccurate outlier predictions suggests that the MLP is more likely to make poor predictions when faced with expression states outside the distribution encountered in its training data, a phenomenon termed “catastrophic forgetting” (French, 1999). To illustrate these MSE values in terms of the simulated expression levels, we take a closer look at a particular cell in Figure 2C. We selected this example as an approximately average performance by both the neural ODE and MLP. Here the MLP maintains a MSE under 0.02 until time t = 8, after which the predictions are inaccurate (median MSE of 0.42 over time points). The neural ODE, in contrast, is a better, though far from perfect fit to the simulated data (median MSE of 0.054). In some cases, the neural ODE predictions demonstrate a closer fit to the data, with median MSE values as low as 0.017 across fifty time points (Supplemental Figure 1). However, we observe some poor fits with the neural ODE as well, producing median MSE values as high as 1.58 (Supplemental Data).

In order to understand why some neural ODE solutions perform substantially better than other solutions, we examined the impact of different gradient descent initializations. Due to the recursive application of the neural ODE, the random seed used to initialize stochastic gradient descent impacts predictions substantially. Even with the exact same training data, neural ODEs with different initializations can yield highly divergent predictions after fifty time points (Figure 2D). We observe that the predictions at time t = 1 are very similar from differently initialized networks, as we would expect, given that the networks are trained on the exact same data. However, because stochastic gradient descent can find many local minima, the weights are somewhat different. When making recursive predictions with the network, these differences in the weights compound, which often leads to very different predictions at later time points (Figure 2D). We observe that some differently initialized neural ODEs perform better than others on a given example, but not uniformly better across all examples (Supplemental Figure 2). This observation suggests that each differently initialized neural ODE may learn slightly different information about how to predict future expression states.

Ensemble based predictions leveraging information across multiple simulations from varied parameters have been shown to improve predictions of complex dynamical systems (Fertig et al., 2007), and are readily adaptable from weather prediction to forecasting biological systems (Kostelich et al., 2011). Therefore, we take an ensemble approach to improve RNAForecaster’s ability to leverage the slightly different information learned by each network and handle variation in prediction accuracy. Using a different random seed to initialize gradient descent for each network, we train multiple neural ODEs and then evaluate the predictions of each, taking the median prediction as the final expression level estimate. This approach substantially outperforms a single network across simulated data sets (Figure 2E) (Supplemental Figure 3). As expected, at time t = 1 there is no significant difference in prediction accuracy, but there is a significant difference by t = 10 (p < 1e-6) and the magnitude of the difference in MSE loss increases with t. Most notably, the ensembles are much less vulnerable to catastrophic forgetting. If one network has extreme outliers in its prediction of gene expression profiles, it will usually be overruled by the others. We find that ten networks are sufficient to achieve most of the accuracy gains we can achieve through ensembling. Twenty-five networks yields no significant improvement over ten networks at any of the fifty time points, though the average MSE across simulations is slightly lower. Using ensembling, RNAForecaster is able to maintain an average log MSE below 0.7 for up to 200 time points (Supplemental Figure 4). A potential concern with ensembling neural networks is the increase in computational costs. However, RNAForecaster remains efficient even on relatively large single-cell datasets by running its computations in parallel or, ideally, on a GPU, which dramatically decreases time and memory costs (Supplemental Figure 5).

To determine the limits over which RNAForecaster can generalize robustly outside its training data, we simulated a single-cell data set using the bifurcating cell lineage gene regulatory network proposed by the authors of BoolODE (Pratapa et al., 2020). In the simulation, the cells progress in a single lineage for about 375 simulated time points, after which they bifurcate into two distinct lineages (Figure 3A). We trained RNAForecaster on time points 365 and 366 across 2000 cells to determine whether training immediately before the bifurcation provided sufficient information for RNAForecaster to predict cell lineage after the bifurcation. We find that RNAForecaster can differentiate the two lineages over 100 predicted time points (Figure 3B) despite being trained before the bifurcation. We compared the MSE on these 100 time points against the median MSE across the random simulations described above. The MSE was lower than the median level on 42 of 100 time points, indicating that RNAForecaster performed comparably in this case to the random network simulations. Immediately before the bifurcation, the expression differences in the cells appear sufficient to indicate which lineage a cell will become. This result demonstrates the ability of RNAForecaster to make predictions outside the space of its training data. However, if trained well before the bifurcation, we hypothesized that it should not be possible to reliably predict cell lineage fate on a per cell basis. To test this hypothesis, we trained RNAForecaster at time points 250 and 251, well in advance of the bifurcation, and predicted through the next 200 time points. We find that predictions break down at the bifurcation point, estimating transcriptional states that fall into a new cluster of cells in the UMAP that did not exist in the simulation (Figure 3C-D). The MSE predictions are likewise poor, worse than the random network median on all 200 time points. This result still leaves uncertainty as to whether the predictions of the second model are worse because of the stochastic effects and recursive error that are unavoidable when predicting from further away in time or due to the model being unable to learn key predictive relationships at the earlier time point. To distinguish these possibilities, we applied the second model (trained at the earlier time points) to predict one hundred time points forward from the cells the first model was trained upon. In this simulation, we find that the second model does predict the bifurcation (Figure 3E-F), though slightly less accurately than the first model. This result indicates that the major predictive relationships can be learned throughout the lineage of the cells, but that predictions only remain accurate over a limited time period due to stochastic effects and error propagation.

**Figure 3.**
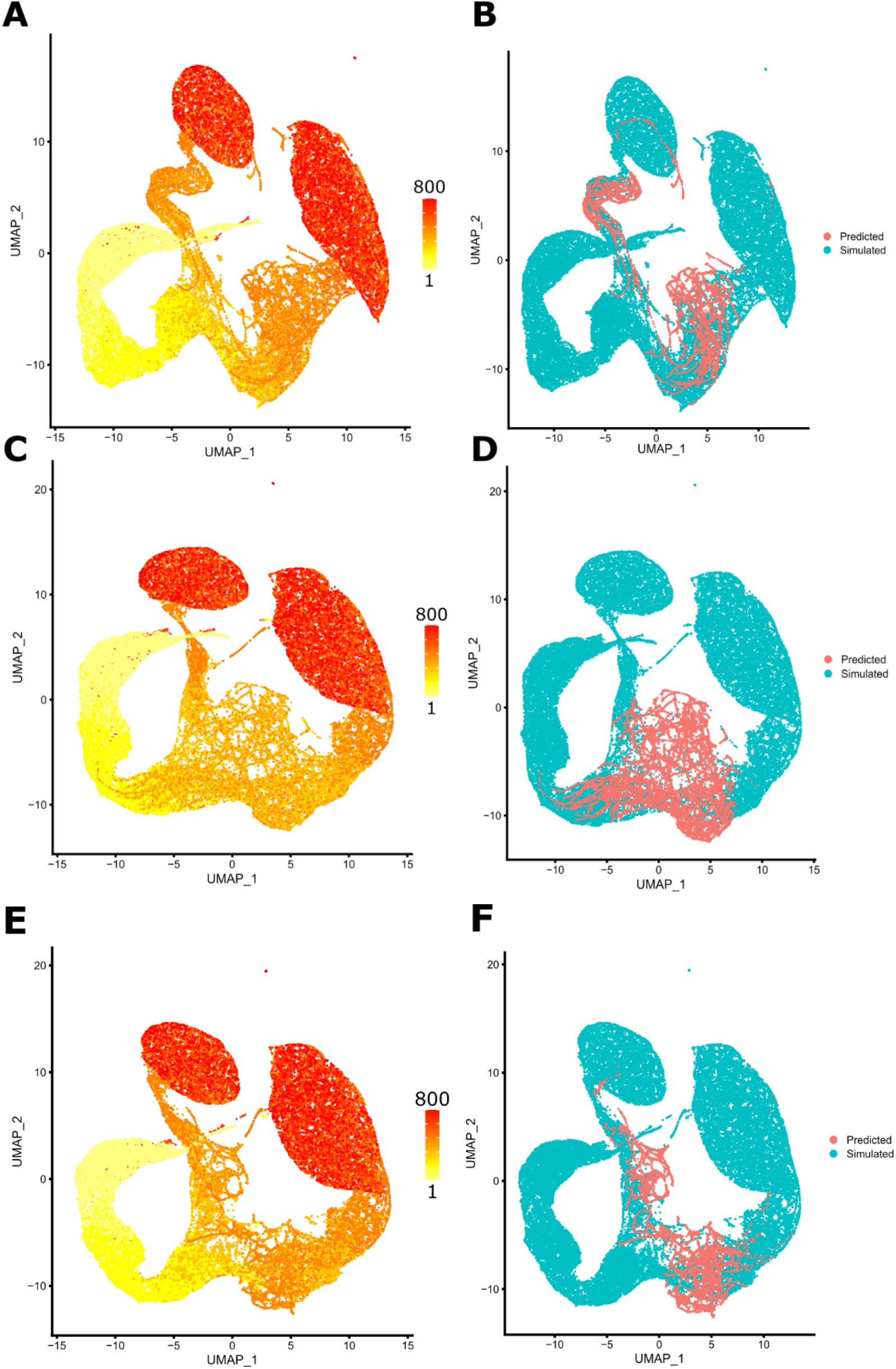
RNAForecaster predicts bifurcation of cells when trained on cells immediately prior to bifurcation **A** UMAP of bifurcating cell simulation across 50 cells from simulated time point 1 to time point 800, plus RNAForecaster’s predictions from time point 366 (just before the bifurcation) through next 100 time points. Colored by time point. **B** UMAP of same cells as in A, but colored by whether cells were from the ground truth simulation or RNAForecaster’s predictions. **C** UMAP of bifurcating cell simulation across 50 cells from simulated time point 1 to time point 800, plus RNAForecaster’s predictions from time point 251 through next 200 time points. Colored by time point. **D** UMAP of same cells as in C, but colored by whether cells were from the ground truth simulation or RNAForecaster’s predictions. **E** UMAP of bifurcationg cell simulation across 50 cells from simulated time point 1 to time point 800, plus RNAForecaster’s predictions using the model from C and D, predicted from time point 366 (just before the bifurcation) through the next 100 time points. Colored by time point. **F** UMAP of same cells as in E, but colored by whether cells were from the ground truth simulation or RNAForecaster’s predictions.

We additionally attempted to use a lower dimensional embedding of the data as input to the neural ODE to determine if this would help with the problem of noise in inherently stochastic data. A variational autoencoder (VAE) has been used with neural network models in other genomics analysis methods for this purpose (Lotfollahi et al., 2019), (Yu and Welch, 2022), (Chen et al., 2022). To test this approach, we trained a VAE on the simulated data from the first two time points. Then, the lower dimensional embedding produced by the encoder part of the network was input to the neural ODE network. The output from the neural ODE was decoded by the VAE back into gene expression levels for each gene. We used an ensemble of ten neural ODEs to compare this approach to an ensemble of ten neural ODEs trained on the individual genes. We found that the VAE approach had significantly higher MSE at all time points than the by-gene RNAForecaster model (p < 1e-7) (Supplemental Figure 6). The difference in performance is largest at early time points and becomes smaller as time goes on. This result suggests that the loss of information from the encoding of the data limits the ability to estimate each gene’s expression level accurately at future time points, especially in the early time points when more accurate predictions are possible.

### RNAForecaster can predict gene expression states beyond the space of the cell states used as input

One distinguishing ability of RNAForecaster from trajectory inference single-cell algorithms is that it can predict gene expression values outside of the space in a two-dimensional embedding occupied by the input data. To demonstrate this feature, we modified the BoolODE simulation to introduce a knock out (KO) of a single gene after one hundred simulated time points, after which the simulation continues with that gene’s value set to zero. The simulated knock out of a single gene further introduces changes in the expression values of all other genes over time, leading to a divergent cluster of cells in UMAP space relative to the simulated cells from earlier time points (Figure 4A).

**Figure 4.**
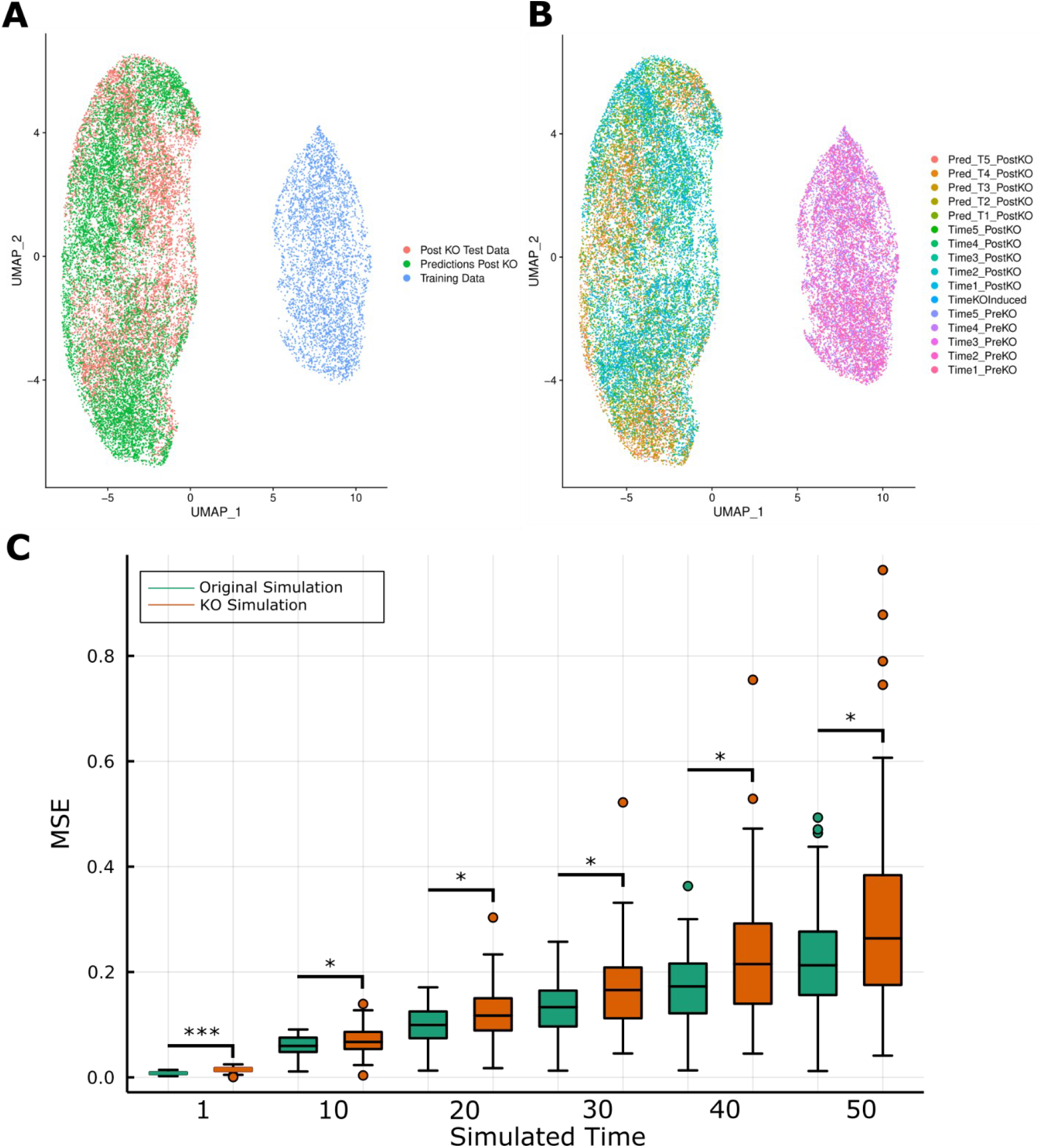
RNAForecaster can predict the impact of a gene KO that moves cell expression outside the input space **A** A UMAP embedding of the training data from one simulation provided to RNAForecaster alongside the simulated data after a gene KO and RNAForecaster’s estimations of expression states after KO. **B** UMAP from same simulation as A, labeled by time point and whether a cell was from the pre-KO simulations, post-KO simulation, or post-KO RNAForecaster prediction. **C** Boxplot comparing the MSE loss from the ten network ensembles shown previously in Figure 2 and the MSE loss from ten network ensembles onto simulated KO data, where the same gene networks were used to generate the simulations in both cases. * p < 0.01 ** p < 1e-6 *** p<1e-10

In order to determine whether RNAForecaster could predict into this space without being trained on it, we trained an ensemble of ten neural ODEs based on only the two time points before the KO simulation began. We then interrogated RNAForecaster’s predictions of future gene expression profiles at time points after the KO occurred. The predicted expression profiles cluster distinctly with the KO simulated data, despite being trained on none of these cells (Figure 4A-B), illustrating the ability of RNAForecaster to make accurate predictions outside the input space it was trained on. Across each of the simulated data sets used to evaluate the ensemble network performance, we simulated a data set from the same regulatory network with a gene KO. We find that RNAForecaster, using an ensemble of ten networks, is capable of producing comparably accurate predictions over fifty time points to those it made on the simulations that did not introduce a KO. We observe a small but statistically significant decrease in prediction accuracy at all time points (the mean difference is less than 0.05 MSE for time points 1 to 30 and less than 0.1 MSE for all time points) (Figure 4C). A loss in predictive accuracy from the distributional shift a KO causes is expected, but the small size of the difference demonstrates that RNAForecaster is able to produce accurate predictions for most simulated KOs through 50 time points.

### RNAForecaster predicts the direction of cell cycle related transcriptomic changes over 72 hours from metabolic labeled scEU-seq data

In order to perform the recursive predictions that allow RNAForecaster to make predictions into the future, we need input data that can approximate the t = 0 and t = 1 matrices we used with simulated data. Critically, these count matrices must contain the two time points from the same cell. Metabolic labeling single-cell RNA-seq is the method we use to accomplish this. With metabolic labeling protocols (such as the scEU-seq protocol (Battich et al., 2020) we will use here) cells are labeled with 4sU modified uridine for a specified time period (Figure 5A). The cells are then harvested for single-cell RNA-seq. The 4sU labeled transcripts can be identified as those that were produced within the labeling period. The other sequenced transcripts were produced before the labeling began. This provides the temporal information we need to train RNAForecaster. The input matrix for the input layer consists of the unlabeled counts matrix plus estimated degraded transcripts. Metabolic labeling data allows for an estimate of the degradation rate in real time, as described by (Qiu et al., 2022). This count matrix is able to represent the total transcripts in the cell at time t = 0. The total counts at t = 1 are then provided by the unlabeled + labeled counts together (Figure 5A).

**Figure 5.**
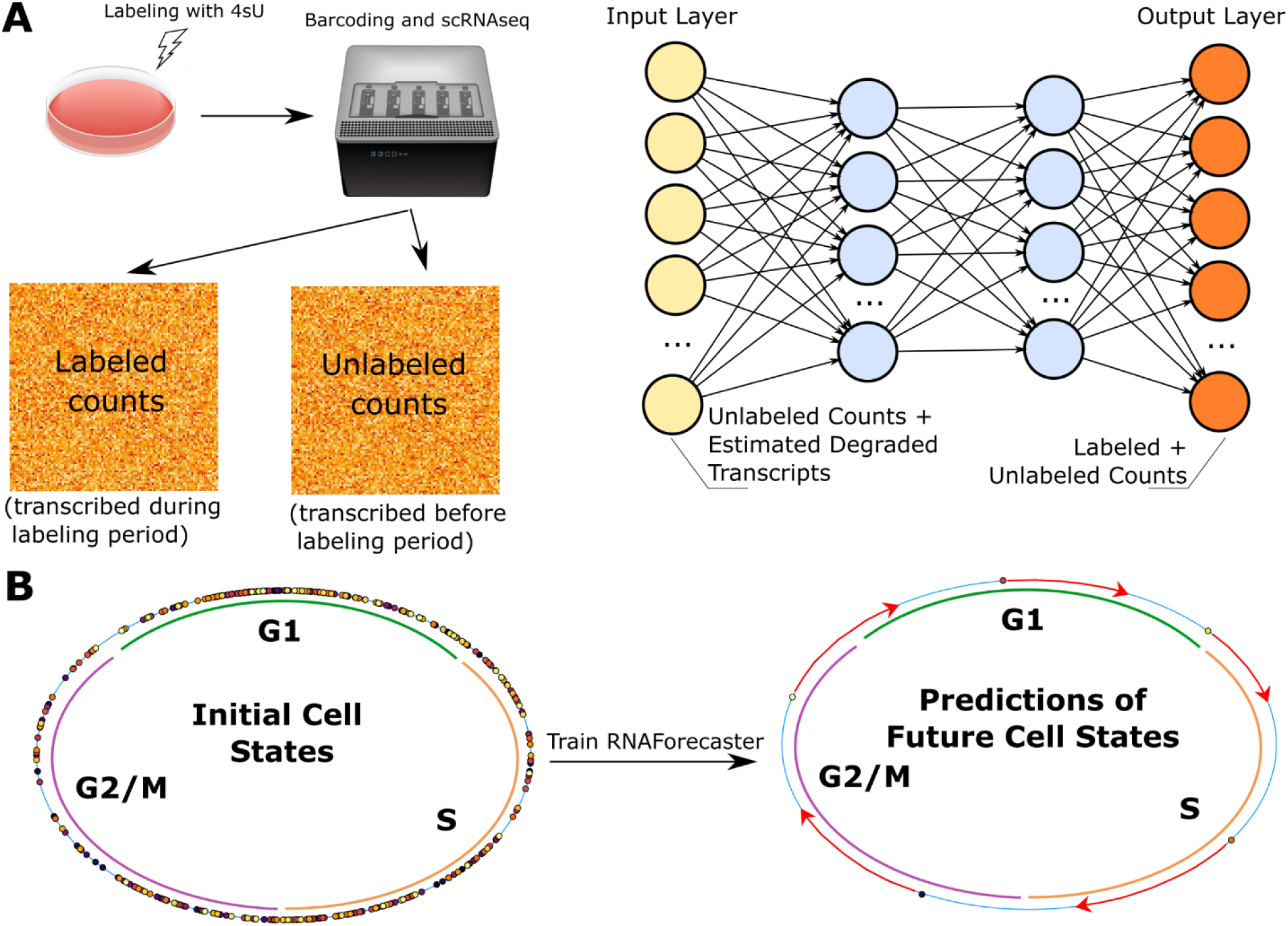
Application of RNAForecaster to metabolic labeling single cell expression data **A** Left is a diagram of the basic concept behind metabolic labeling protocols such as scEU-seq. On the right is a diagram illustrating how the output from metabolic labeling protocols is input to the RNAForecaster neural network. **B** Diagram showing the tricycle cell cycle scores of each one hour labeled cell from the Battich, et al (2020) scEU-seq retinal epithelium cell cycle data set. After these cells are used to train RNAForecaster, the future expression states of each cell can be predicted. These expression states can likewise be scored for cell cycle prediction and we can validate the predictions on whether they generally follow the expected trajectory of the cell cycle.

Now that we have a framework that allows us to train RNAForecaster on a biological data sets, we need a method for assessing its performance. We cannot get a series of expression levels in the same cell, preventing a direct assessment of per gene error over time. However, we can train RNAForecaster in a context where we know the general future expression path the cells should take, such as the cell cycle. To validate the method, we can test if RNAForecaster is able to predict the transcriptomic changes that are required for cell cycle progression. For this validation, we employed a scEU-seq data set from immortalized human retinal pigment epithelium (RPE) cells, published by (Battich et al., 2020). RNAForecaster was trained on the 405 cells in the data with a one hour labeling period, using a ten network ensemble. Once trained, RNAForecaster was used to predict the future expression levels in each cell for 72 hours. To score each cell’s position in the cell cycle, we used tricycle, an R package that projects gene expression data onto an embedding of well characterized cycling cells to create a continuous score for cell cycle position (Zheng et al., 2021). These scores range from 0 to 2π, allowing cell cycle scores to be visualized on a circle (Figure 5B). The expression level predictions of RNAForecaster are then likewise scored by tricycle. The degree to which they are ordered from 0 to 2π over time is then assessed to estimate the degree to which RNAForecaster accurately predicts cell cycle progression.

The tricycle scores are highly ordered with respect to time through 72 hours of predictions in most cells (Figure 6A). For all 72 hours, the predictions are significantly more ordered than randomly generated scores (p < 1e-16). The ordering of these scores relative to random was further checked by generating 10 million random sets of tricycle scores, none of which achieved an ordering score equal to or greater than the median order score from the RNAForecaster predictions. Within the RNAForecaster predictions, the order of the scores decreased significantly each day (p < 0.001), indicating the fidelity to the cell cycle and general quality of predictions decreased the further predictions were into the future, as expected due to error propagation with recursive prediction. A challenge we dealt with in this data was a tendency of the neural ODE to eventually start predicting extremely high transcript counts (Figure 6B). This likely results as an example of catastrophe when the network encounters input that is sufficiently dissimilar to what it was trained on. Then, in an example of positive feedback in the predictions, the predicted gene expression levels begin to go towards infinity. In order to control these extreme values in the prediction model, we set a realistic prior on the upper bound of expression values given the expression distribution observed in the training data. Enforcing these priors yields predictions that have similar median total counts/cell as in the scEU-seq data, even after 72 recursive predictions (Figure 6B). These maximum counts priors also improve performance on cell cycle ordering scores by a small but statistically significant amount (p < 0.05) (Figure 6C).

**Figure 6.**
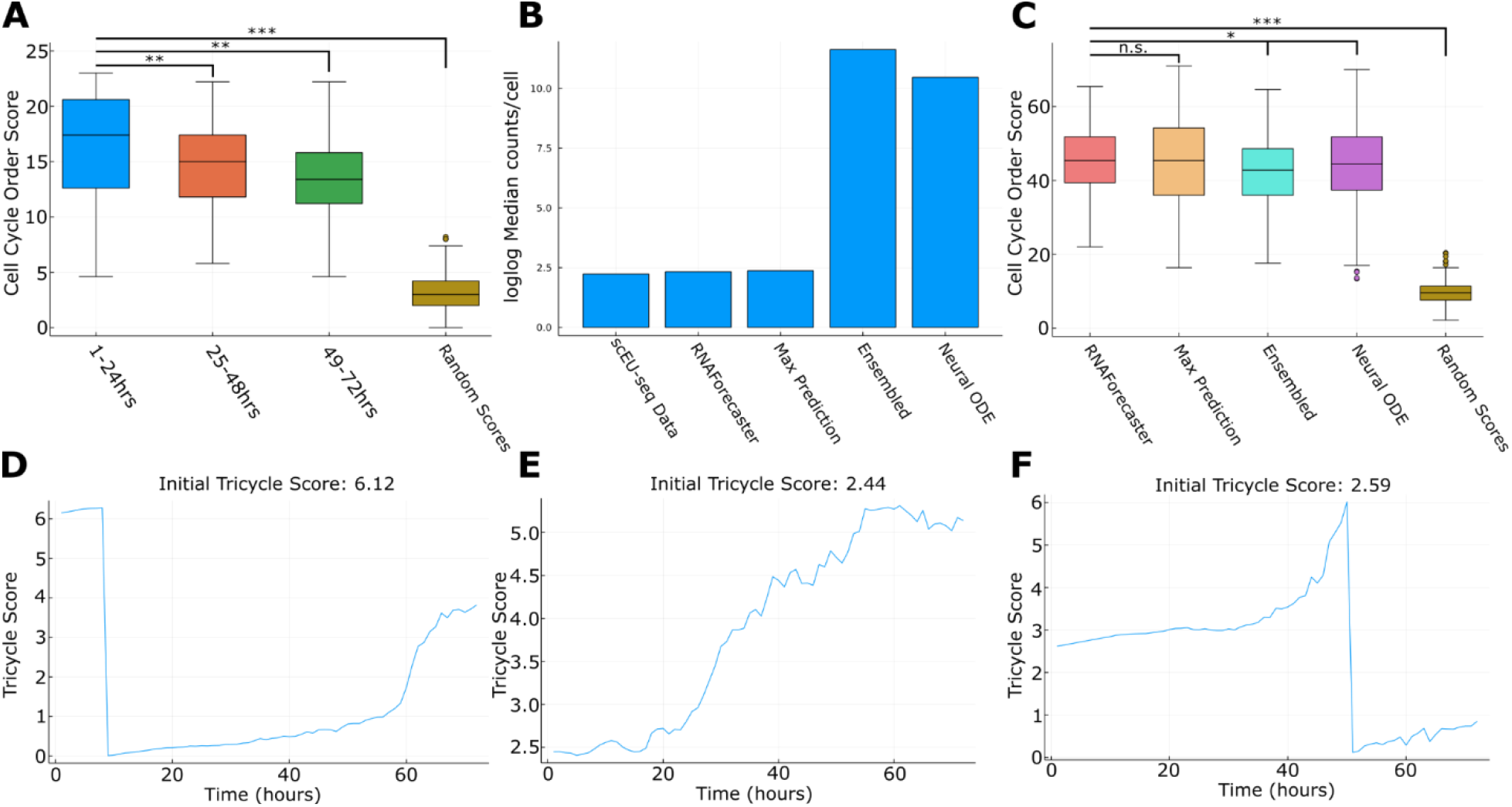
Performance of RNAForecaster at forecasting the cell cycle **A** Boxplot of a metric describing the order of the tricycle scores made using RNAForecaster’s predictions. A higher score indicates the scores were more aligned with the order of the cell cycle. Compared to the metric when applied to randomly generated tricycle scores. **B** Barplot of the log-log median total counts per cell in the scEU-seq data set vs the output of different neural ODE implementations at the 72 hour prediction. **C** Boxplot of the tricycle score order metric for the neural ODE implementations shown in B. **D-F** Examples of tricycle scores on the RNAForecaster predictions in three cells. * p < 0.05 ** p < 0.001 *** p < 1e-16

Tricycle is used to score where RNAForecaster’s predictions are in the cell cycle at each hour over the three day period, allowing us to determine the rate at which the predictions are moving through the cell cycle. While the order of scores strongly reflects the cell cycle, the rate of progression predicted by RNAForecaster is much slower than expected. Immortalized RPE cells usually replicate about once every 24 hours, and the RNAForecaster predictions proceed much more slowly (Figure 6D-F) (Supplemental Figure 6). Cells generally are predicted to progress steadily through the cell cycle, tracking the order of the cell cycle, but falling well short of the expected three completed cycles. This observation indicates that RNAForecaster learns the standard movement of cell cycle related genes, but is unable to recognize rarer regulatory events that lead to large changes in expression over shorter time periods. A relatively small number of cells were available for training RNAForecaster (405) which may contribute to this issue. Particularly for less common events, a larger data set could allow RNAForecaster to more adequately learn to model these gene expression dynamics. Alternatively, predicting shifts that are less frequently observed in the training data may be a weakness of the RNAForecaster’s neural network architecture. Despite this limitation, accurately predicting the order of changes in cell cycle related genes across the different cell cycle stages using only a relatively small training set demonstrates the ability of RNAForecaster to estimate future expression states in single cells.

## Discussion

RNAForecaster is a tool for generalizing temporal relationships in single cell transcriptomic data. Through a neural ODE, we attempt to learn activation functions that predict the expression level of a gene in terms of the previous expression levels of other genes. We demonstrate that it is possible to forecast future expression states in single-cell data that have a temporal dimension. The accuracy of the predicted gene expression states depends on the time period over which RNAForecaster is applied, with a reasonable degree of accuracy over short to intermediate time periods. RNAForecaster can thus provide valuable insight into the dynamics of transcription and transcriptional regulation over time. Through simulated data, we demonstrate that KOs and bifurcations can be predicted if the method can predict starting at time points shortly before the event occurs. In order to better capture the relationships between genes that allow prediction of their future expression states, it would be ideal to train the RNAForecaster network on more time points within the same single cell. While this is currently unavailable across at scale, recent techniques allow imaging of a small number of genes in cells over time (Cawte et al., 2020), (Wang et al., 2022). As these methods improve, RNAForecaster could be trained with longer time series, likely improving its accuracy and the span of time over which it can make accurate predictions. Altogether, future research should evaluate the limits of predictability using this model over diverse timescales and biological conditions.

The reliance on ODEs in our framework results in estimates of smooth temporal trajectories of gene expression. However, it is important to consider the predictions RNAForecaster produces in the context of the biology of transcription. RNA transcription occurs in bursts, and thus appears stochastic (Tunnacliffe and Chubb, 2020). Therefore, RNAForecaster’s predictions should not be interpreted as an estimate of the exact counts in a cell, since this is not precisely predictable. Rather, RNAForecaster should be thought of as estimating the expected value of the distribution of a gene’s counts in a cell.

In simulated data, we observed that significant gains in prediction accuracy could be attained by using an ensemble of networks to forecast future expression states instead of a single network. Given the recursive application of the network to make future predictions, the impact of small differences in the network weights can lead to large differences in predictions when two different initializations of stochastic gradient descent are used. The slightly different local minima found by these different gradient descent initializations often had strengths and weaknesses in their ability to generalize to expression levels outside their training data. We attempted to combine these strengths through a simple ensemble approach where we use the median predictions of the networks. However, even with the ensembling approach, prediction accuracy decreased substantially over time in the simulated data as error propagated and predictions trended further outside the domain the neural network was trained in. At the same time, catastrophically poor predictions occurred at a much lower rate, which does increase the time scale in which RNAForecaster is applicable. The downside of this approach is the increased computational resources required. However, training ten networks using a GPU is faster on single cell transcriptomics sized data sets than training one on a CPU (Supplemental Figure 5), which makes GPU training highly preferred even without ensembling.

Using metabolic labeling scRNA-seq protocols, RNAForecaster can make recursive predictions about future transcriptomic states. These protocols are currently largely confined to *in vitro* studies, which limits the application of RNAForecaster in many *in vivo* contexts. *In vitro*, RNAForecaster was capable of accurately predicting the general direction of cell cycle related expression changes over 72 hours. However, the model failed to recapitulate the speed of cell cycle progress. This may reflect an inability of the model to predict cell cycle checkpoints and the ensuing transitions between stages, or some other rare regulatory event in the cell cycle. A larger training set might allow RNAForecaster to better capture these less common events, given that the training data contained only 405 cells. However, it may be the case that the current structure of RNAForecaster lacks the capability to handle these sorts of exceptions to the transcriptomic changes it sees in most cells. Future work integrating attention based architecture (Vaswani et al., 2017) into RNAForecaster could potentially allow the model to differentiate, for example, the expression changes within a cell cycle stage from the changes at the end of the stage after checkpoints are passed.

We observed that the neural ODE tended to make extremely high predictions of expression values after the previous predictions had departed sufficiently from its training data. To handle this we enforced maximum expression level predictions for each gene based on the observed data, which constrained the model to obtain realistic expression levels on both an individual gene and overall cell level. These maximum expression levels can be justified from a Bayesian perspective, where we assign very low probability to seeing expression levels of a gene that are higher than a certain point. There is a difficult balance, however, between preventing unrealistic expression levels and removing valuable signal from the predictions. Preventing these kinds of extreme, unrealistic values is a major challenge termed as “catastrophe” in the field of machine learning, (French, 1999), and it may be further exacerbated for RNAForecaster due to the high dimensionality of the data and the recursive predictions required. The use of network ensembles helps alleviate this tendency to some extent. However, future work may be needed to teach the neural network about the prior probability for gene expression levels outside the range normally observed in cells, rather than having to enforce this prior after the fact.

Whereas many predictive methods, including notably Dynamo (Qiu et al., 2022), can estimate cellular states from RNA velocity vector fields estimated through splicing or metabolic labeling, RNAForecaster relies on temporal single-cell transcriptomics data tracing an individual cell currently enabled uniquely with metabolic labeling data (Qiu et al., 2022). Whereas these current methods aim to predict cellular states captured in the training data, RNAForecaster instead attempts to generalize its predictions to the full space of possible expression states. This formulation uniquely allows RNAForecaster to estimate the impact of perturbations that shift the expression state into part of the space not observed in the input, as well as future developmental or evolutionary states not captured in the input data. Another important difference between Dynamo and RNAForecaster is that Dynamo requires its input data to be smoothed using k-nearest neighbors averaging in order to compute RNA velocity (Qiu et al., 2022). This procedure essentially averages the cells that are close together in expression space, which may introduce some distortions or remove important variation (Gorin et al., 2022).

Several other methods have been proposed with the goal of predicting single-cell responses to perturbations. PerturbNet trains a generative neural network using perturbation single-cell data sets such as Perturb-seq (Dixit et al., 2016) to predict responses to genetic knockouts or knockdowns (Yu and Welch, 2022). This approach differs from RNAForecaster by focusing on learning from specific perturbations rather than learning general temporal relationships between genes. The scGen method also attempts to predict single-cell perturbation responses, using a combination of a variational autoencoder and a deep generative network to project what it has learned in its training data into unseen cell states (Lotfollahi et al., 2019). While RNAForecaster attempts to learn the predictive relationships between genes over time, scGen attempts to learn how cell states shift under perturbation using similar perturbations in similar cells. These approaches can yield valuable insight into future expression states after a perturbation and the correct choice of method for a particular use case will often depend on the particulars of a problem and the type of data that is most readily available.

The gold standard that many computational methods aspire to is inference of mechanistic interactions from high-throughput biological data sets. One advantage of neural ODEs is that they can yield greater interpretability than other neural network formulations (Chen et al., 2018). This feature may allow the relationships between genes that RNAForecaster learns to be interrogated, which could potentially yield mechanistic insight. The accuracy of gene network inference methods suggests that the high degree of correlation between genes makes prediction much easier and more robust than causal inference (Pratapa et al., 2020). However, extending RNAForecaster and other methods from the prediction of future gene expression states to mechanistic, molecular networks remains an important area of future research.

RNAForecaster demonstrates that future states in transcriptomic data with a temporal dimension can be estimable, even outside the expression space of the input data. As single-cell and machine learning technology improves, it may be possible to extend this capability to accurately predict counterfactuals regarding future cell states based on a variety of cellular factors. This could be used to predict the response of diseased cells to perturbations, potentially informing treatment options on a general or personalized level. Extending these techniques to personalized predictions of the effects of perturbations and therapies may enable predictive biology and medicine approaches (Fertig et al. 2021), (Stein-O’Brien, Ainsile, and Fertig 2021) and require new methods to quantify the limits of predictability of therapeutic outcomes across disease systems.

## Methods

### RNA Forecaster

#### Required Input Data and Preprocessing

RNAForecaster primarily requires two normalized single-cell RNA counts matrices as input. These counts should be from adjacent time points, such that the labels time t=0 and time t=1 can be reasonably applied and the cells in each matrix are identical. The main preprocessing steps needed are sparsity filtering and log normalization. Including genes that have high proportions of zeroes (default greater than 98%) can cause problems with gradient descent, and thus these genes must be removed. The only normalization applied is a log1p transform. In addition, filtering to highly variable genes is strongly recommended, and was performed for all biological data sets used.

When using metabolic labeling data, there is an additional preprocessing step to account for transcripts that degraded during the labeling period. The degradation rate is calculated using the slope between the labeled and total counts as described by (Qiu et al., 2022). A linear regression is fit between the two count matrices and the degradation rate is calculated as *-log(1-slope)* which estimates the number of transcripts degraded per labeling period. We then estimate the total counts at the beginning of the labeling period by adding each gene’s degradation rate to the unlabeled count matrix. The resulting matrix becomes the time t=0 input matrix.

#### Neural ODE Training

By default, the input data matrices are divided into training and validation sets (default 80-20 split). The default number of nodes in the hidden layer is twice the number of nodes in the input layer to allow for interactions between genes. A neural ODE (Chen et al., 2018) is then trained using Flux.jl and DiffEqFlux.jl, using the Tsit5 ODE solver and a default error tolerance of 1e-3. Training occurs for a default of 10 epochs using a default learning rate of 0.005. The loss function is calculated as the mean squared error between the output nodes and each gene expression level in the time t=1 matrix.

We provide the option to check network stability on recursive predictions at this stage. Recursive predictions are made for a user-defined number of steps, checking on each step whether any expression levels are higher than any plausible level. If this is observed, the training process is restarted. These stability checks can prevent the frequency of catastrophe in network predictions outside the training distribution (Supplemental Figure 7). With ensembling, these stability checks are largely unnecessary, but they provide an alternative for data sets where the network is less prone to catastrophe.

Creating an ensemble of networks simply repeats this process using a different random seed to initialize stochastic gradient descent for each network. The default number of networks trained is ten.

#### Recursive Predictions of Future Expression Levels

The process of estimating future expression requires the input of a trained neural ODE (or ensemble of them) and a set of initial expression states to predict from. These expression states are fed into the input nodes of the neural ODE and then the output is recorded and fed back into the input nodes, allowing for recursive predictions forward in time. Some prior knowledge and assumptions are enforced on the predictions by default. All predictions must be non-negative, as this is a constant characteristic of gene expression data. Additionally, expression level predictions that are higher than an allowed maximum are set to the maximum value (by default two times the maximum observed in the training data, in log space).

When estimating expression levels using an ensemble of networks, the above process is performed for each network and the median prediction is used.

#### Variational Autoencoder Input Option

We tested using a lower dimensional representation of the single cell transcriptomics data as input with a VAE. The VAE was trained on the first two time points of the data, the same portion that would be used to train the neural ODE. The encoder was then applied to the t=0 data and the lower dimensional matrix was used as input to the neural ODE, which was trained on the lower dimensional encoding of the t=1 matrix. The output lower dimensional representation was then decoded with the VAE back into individual gene expression levels.

### RNAForecaster in simulated single cell transcriptomics data

#### Simulating single cell expression data with BoolODE

BoolODE was designed to simulate single-cell expression data sets on the basis of a network of gene-gene interactions (Pratapa et al., 2020). We generated 117 simulations because we wanted at least one hundred and we were concerned that some random networks might generate strange regulatory behaviors. To generate these 117 different simulations, we first needed 117 different gene-gene networks. These were created as random ten gene networks, where genes could have positive, negative, or no direct relationship with other genes. Each network was input to BoolODE, which simulates 801 time points of expression for 2000 cells in each simulation. Minor changes were made to BoolODE code (see https://github.com/FertigLab/RNAForecasterPaperCode), to generate output for the task of predicting future expression states.

The bifurcation simulated example was produced using the bifurcation gene network and initial conditions set created for and provided by BooODE (Pratapa et al., 2020). We simulated 2000 cells for 801 time points to yield the data used in this work. UMAP visualizations of the data were produced using Seurat version 4.0.1.

To simulate data sets with gene KOs, the simulations were allowed to run for 101 time points, at which point a gene’s expression value was set to zero and was again set to zero on each future iteration, mimicking a KO gene.

#### Applying RNAForecaster to simulated data

We used the t=101 simulated count matrix as our t=0 for input to RNAForecaster, in order to give the simulation time to initialize and stabilize. RNAForecaster was thus trained on t=101 and t=102 for each simulation. Predictions were made for up to 200 time points later, up to t=300. The neural ODE was trained for 10 epochs, with 100 hidden layer nodes, and a learning rate of 0.005.

Ensembles were created using groups of 10 and 25 networks. Networks used 100 hidden layer nodes and stability checks were performed. Simulations where stability checks were not passed on fifteen iterations were excluded, leaving 111 simulations in the final set.

#### Comparison predictions using a feed-forward MLP model

For comparison with RNAForecaster’s predictions in simulated data, we employed a simple five hidden layer, fully connected feed-forward neural network architecture. This MLP model was trained for 10 epochs and a learning rate of 0.005. The network node structure from the input to output layer is as follows: Dense(10,32), Dense(32,64), Dense(64, 100), Dense(100,100), Dense(100,64), Dense(64, 32), Dense(32,10).

### RNAForecaster in scEUseq from hTERT immortalized human retinal epithelium cells

#### Data download and processing

The scEU-seq data set from (Battich et al., 2020) was downloaded using the Dynamo python package to acquire the rpeLabeling.h5ad AnnData file produced by (Qiu et al., 2022).

Genes with more than 98% zero counts in either labeled or unlabeled count matrices were filtered from both matrices. The matrices were additionally filtered to genes by variance to genes in the top quartile. The degradation rate was then calculated using the slope between the labeled and total counts. A linear regression was fit between the two count matrices and the degradation rate was calculated as *-log(1-slope)* which estimates the number of transcripts degraded per labeling period. We then estimate the total counts at the beginning of the labeling period by adding each gene’s degradation rate to the unlabeled count matrix. We then subset to those cells treated with 4sU uridine for 60 minutes, so that the labeling time is equal for all input cells. The resulting matrix becomes the time t=0 input matrix, to be compared to the total counts as the t=1 matrix in RNAForecaster.

#### Training RNAForecaster

RNAForecaster was trained as an ensemble of ten networks, training for 20 epochs on all 405 cells with a 60 minute labeling period. These networks were trained on a Nvidia Titan V GPU using a mini-batch size of 100 and a learning rate of 0.001. All other parameters use the default values.

#### Predicting future expression states and estimating their position in the cell cycle

RNAForecaster predicts the future expression levels in each cell from the total counts matrix into the future each hour for 72 hours. The maximum prediction for each gene is set to 1.2 log fold increase over the maximum value observed in the training data. Predictions were performed on a Nvidia Titan V GPU.

The resulting predictions were scored for their position in the cell cycle using the tricycle R package (Zheng et al., 2021). Tricycle creates a quantitative embedding of the cell cycle from scRNAseq data of cells with known cell cycle positions. This embedding ranges from 0 to 2π to represent the circular nature of the cell cycle. In this range, 0.5π to π is the approximate bounds of S phase, π to 1.75π G2M phase, and 1.75π to 0.25π G1 or G0 phase. This embedding is then projected into a target single cell RNA data set to approximate the cell cycle position of each cell. We applied tricycle to each initial cell state from the scEU-seq data and each expression state predicted by RNAForecaster.

To determine the degree to which the tricycle scores in the RNAForecaster predictions matched the order of the cell cycle, we developed an ordering metric. The predictions made in a cell receive a point in this metric if the score increases (or goes back around from 2π to 0), but does not increase by more than 0.75 in a one hour period. This metric additionally differentiates small decreases in cell cycle score from large ones by giving 0.2 points to decreases of less than 0.25. This is used to differentiate slight variation from substantial incorrect shifts in gene expression prediction. We additionally plotted the scores, both for individual cells as a line plot, and all together on a circle plot.

## Data and Code Accessibility

Simulated data generated for this study is available on Figshare: https://doi.org/10.6084/m9.figshare.20123804 https://doi.org/10.6084/m9.figshare.20425467

The hTERT human retinal epithelium scEU-seq data (Battich et al., 2020) is archived as an AnnData object on Figshare: https://doi.org/10.6084/m9.figshare.20126492 RNAForecaster is freely available as a Julia software package on GitHub: https://github.com/FertigLab/RNAForecaster.jl and archived on Zenodo: https://doi.org/10.5281/zenodo.6773296.

All code used to perform the analyses described in this paper is published on GitHub: https://github.com/FertigLab/RNAForecasterPaperCode.

## Acknowledgements and Funding

This work was supported by U01CA212007 (to E.J.F.), U01CA253403 (to E.J.F.), and K99NS122085 from the BRAIN Initiative in partnership with the National Institute of Neurological Disorders (G.S.O.).

## Supplemental Figures

**Supplemental Figure 1.**
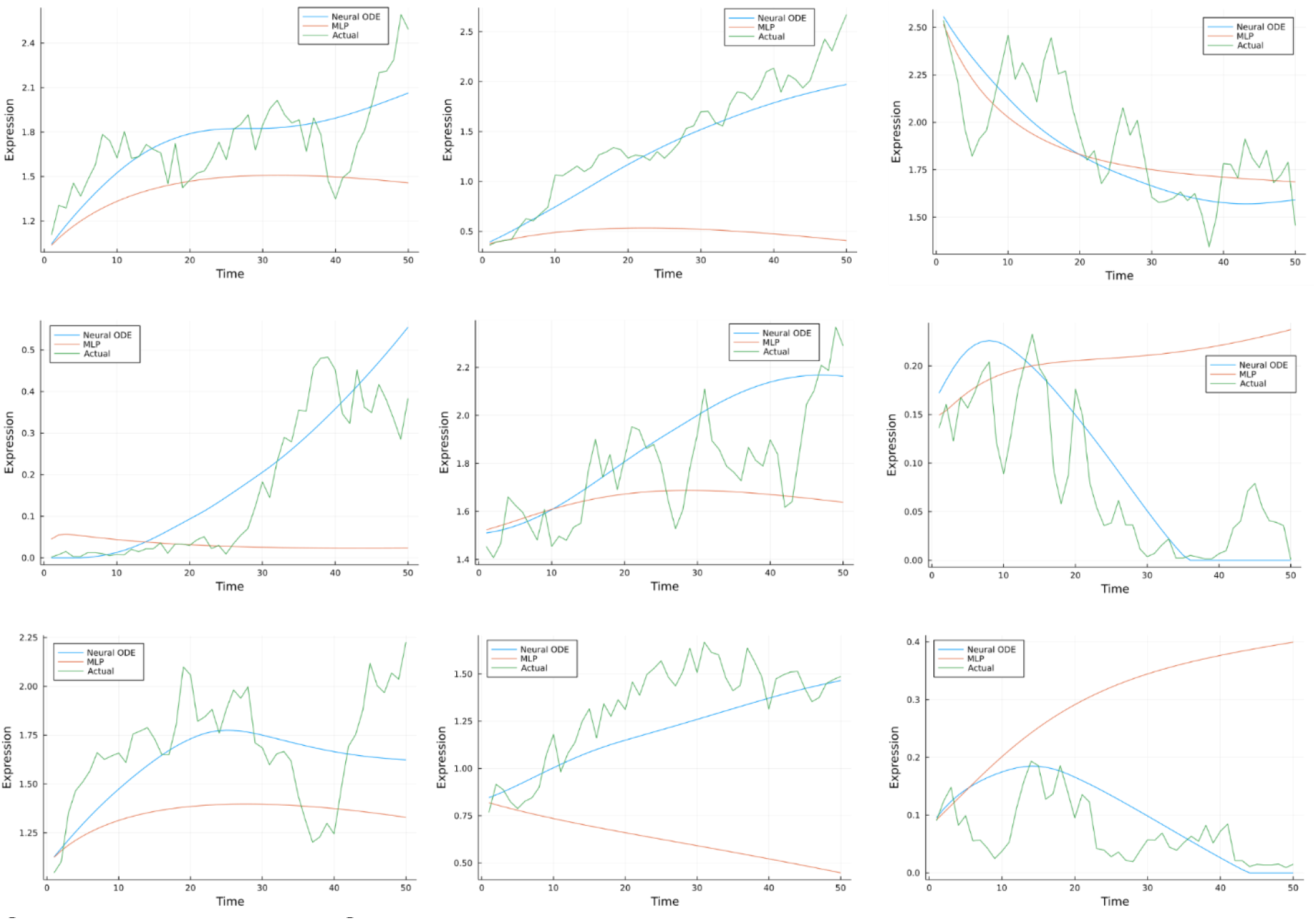
Single cell examples of predictions accuracy over time A set of examples from the simulated data illustrating the fidelity of the single neural ODE network predictions in some cases.

**Supplemental Figure 2.**
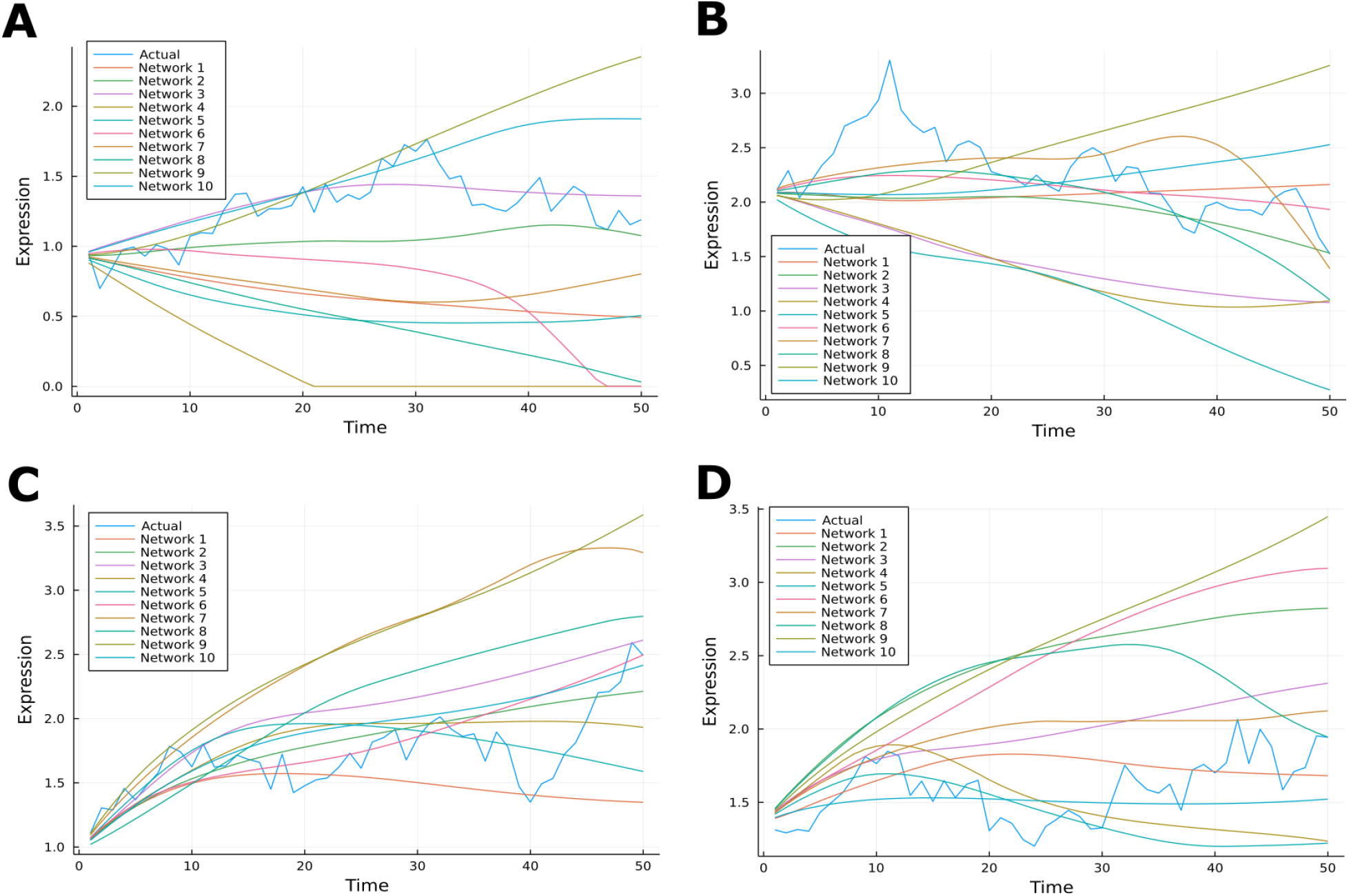
Network accuracy varies by context Figure demonstrating that neural ODEs trained with a different initialization of stochastic gradient descent not only produce different predictions, but the quality of predictions varies between examples. In **A**, Network 3 is highly accurate, but quite inaccurate in **B**. Similarly, while Network 6 is a very good fit in **C**, the fit is very poor in **D**.

**Supplemental Figure 3.**
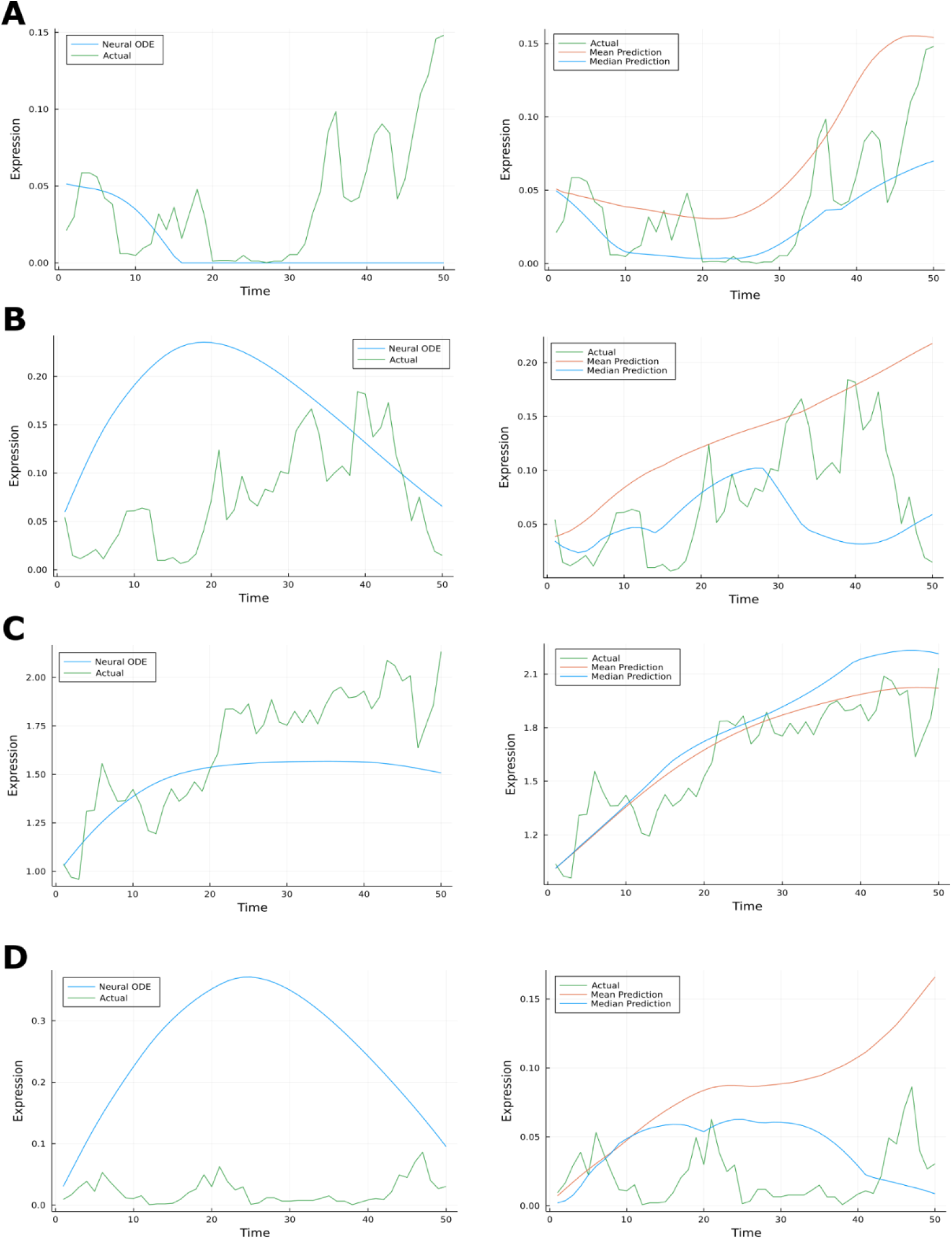
Network ensembling often rescues poor performance on individual cells **A-D** On the left the single neural ODE prediction is shown against the simulated expression level in a single cell. On the right the mean and median prediction of a ten network ensemble is shown for the same cell.

**Supplemental Figure 4.**
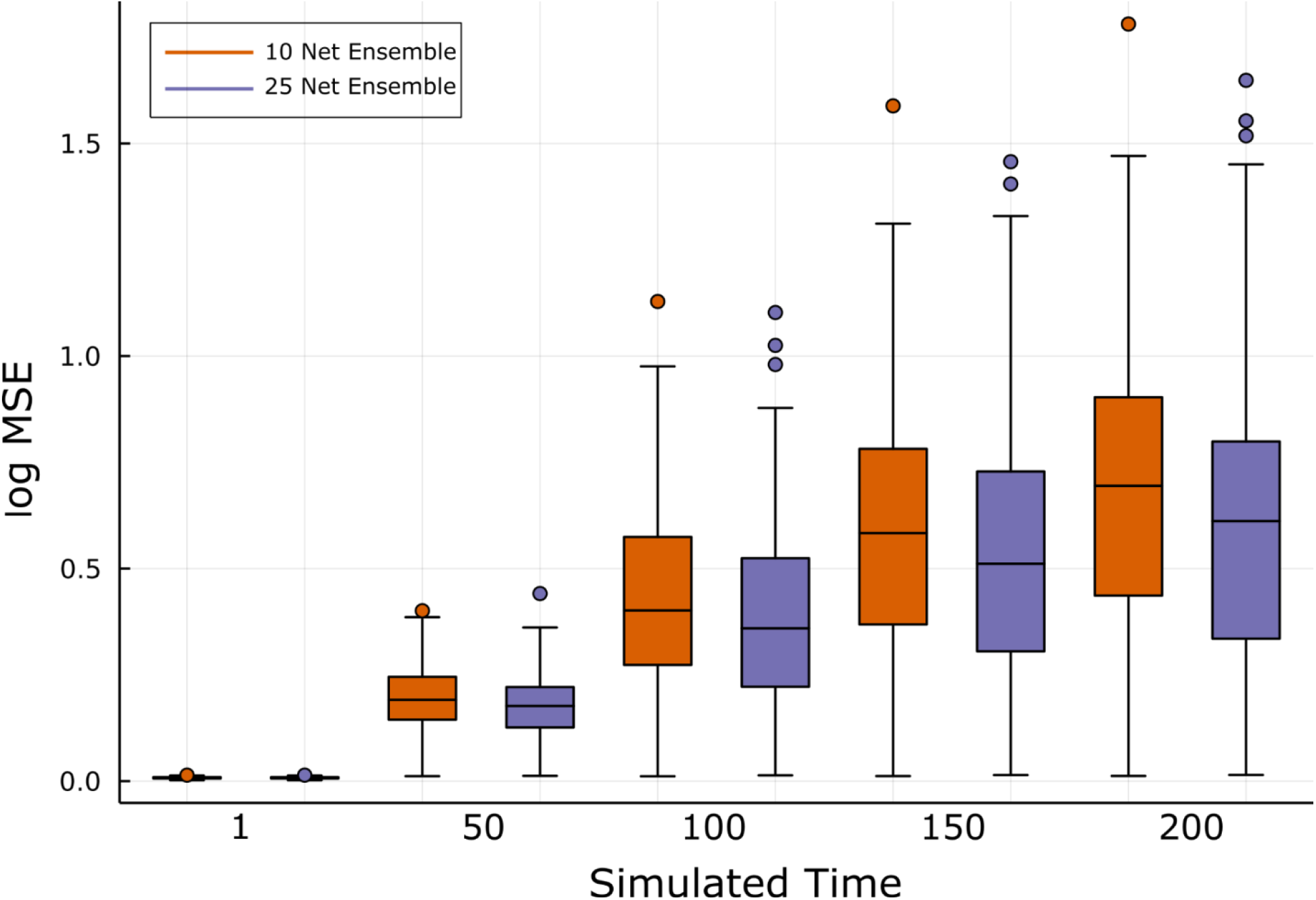
Prediction accuracy of ensembled neural ODEs over 200 time points Log MSE loss from 1 to 200 simulated time points using 10 and 25 network ensembles of neural ODEs.

**Supplemental Figure 5.**
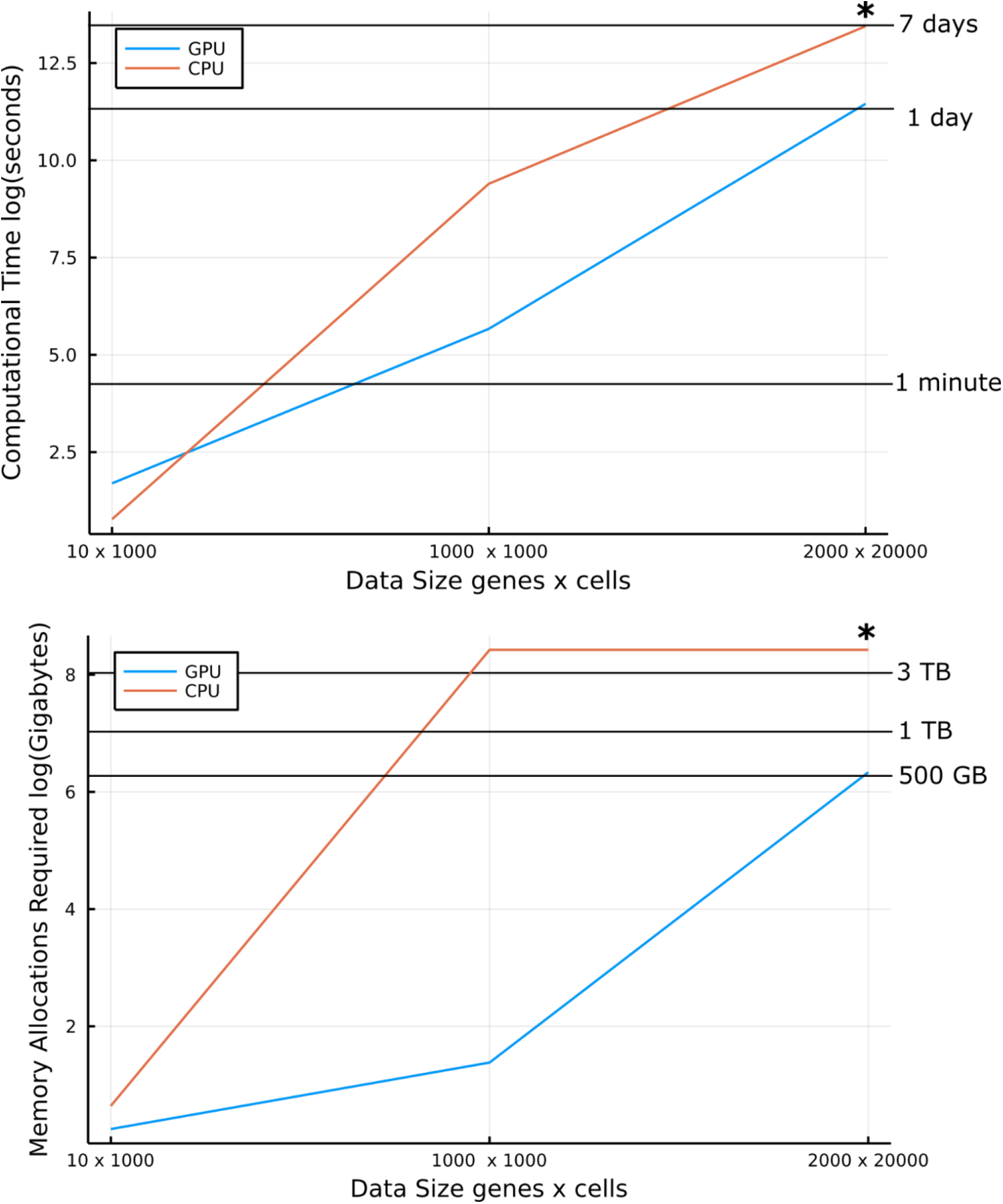
Computational performance of RNAForecaster on a GPU v CPU **A** Time required to train RNAForecaster using GPU vs CPU computation. **B** Memory allocations required to train RNAForecaster using a GPU vs a CPU. * The CPU training for the 2000 gene x 20000 cell data set was stopped after running for 7 days without finishing, so total memory allocations are not reported for the CPU on this data set.

**Supplemental Figure 6.**
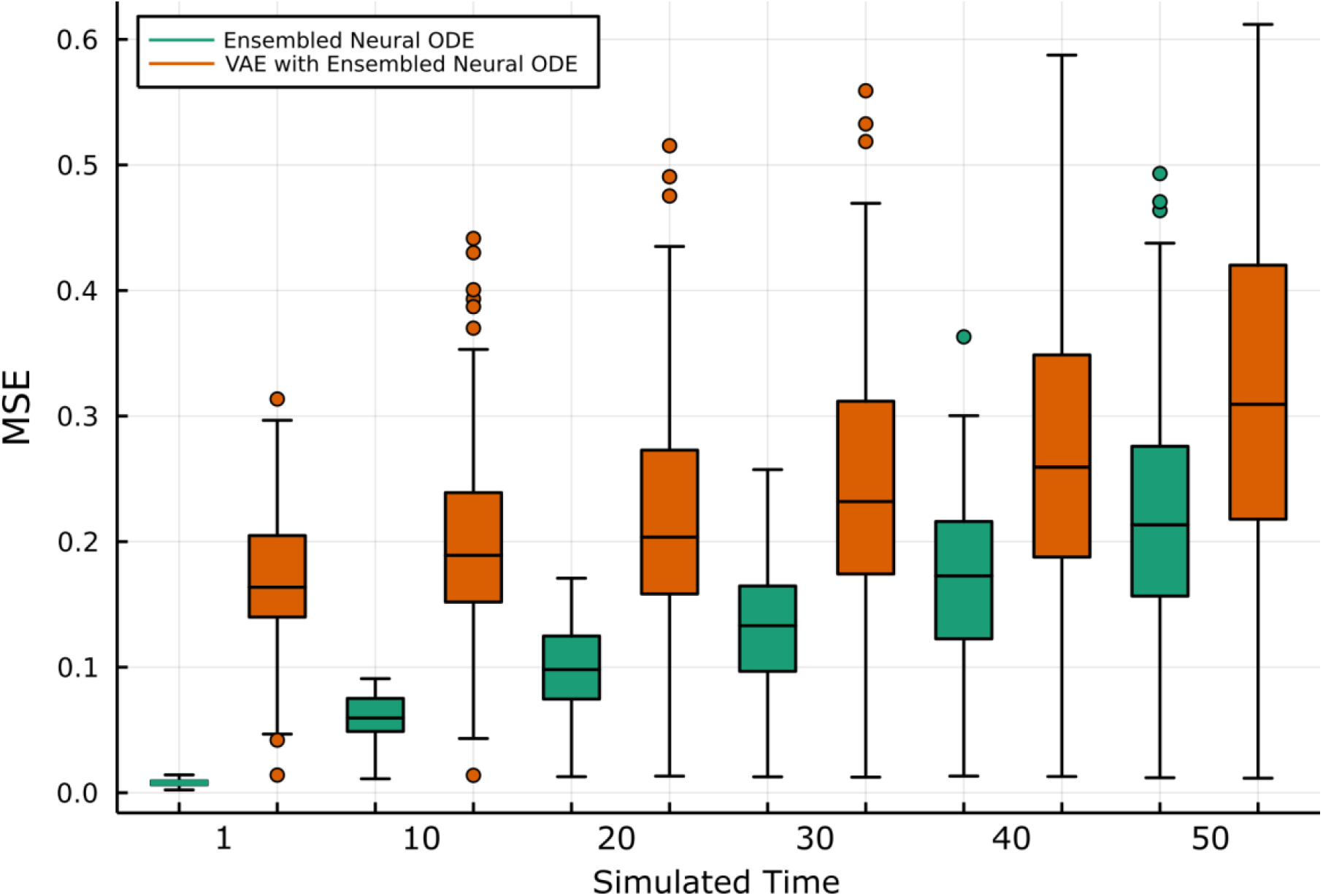
Prediction Accuracy of VAE encoded input vs by-gene input Boxplot across the simulated data sets comparing performance of a neural ODE ensemble where the input was a VAE encoding of the expression data against a neural ODE ensemble where the input was each gene’s expression level.

**Supplemental Figure 7.**
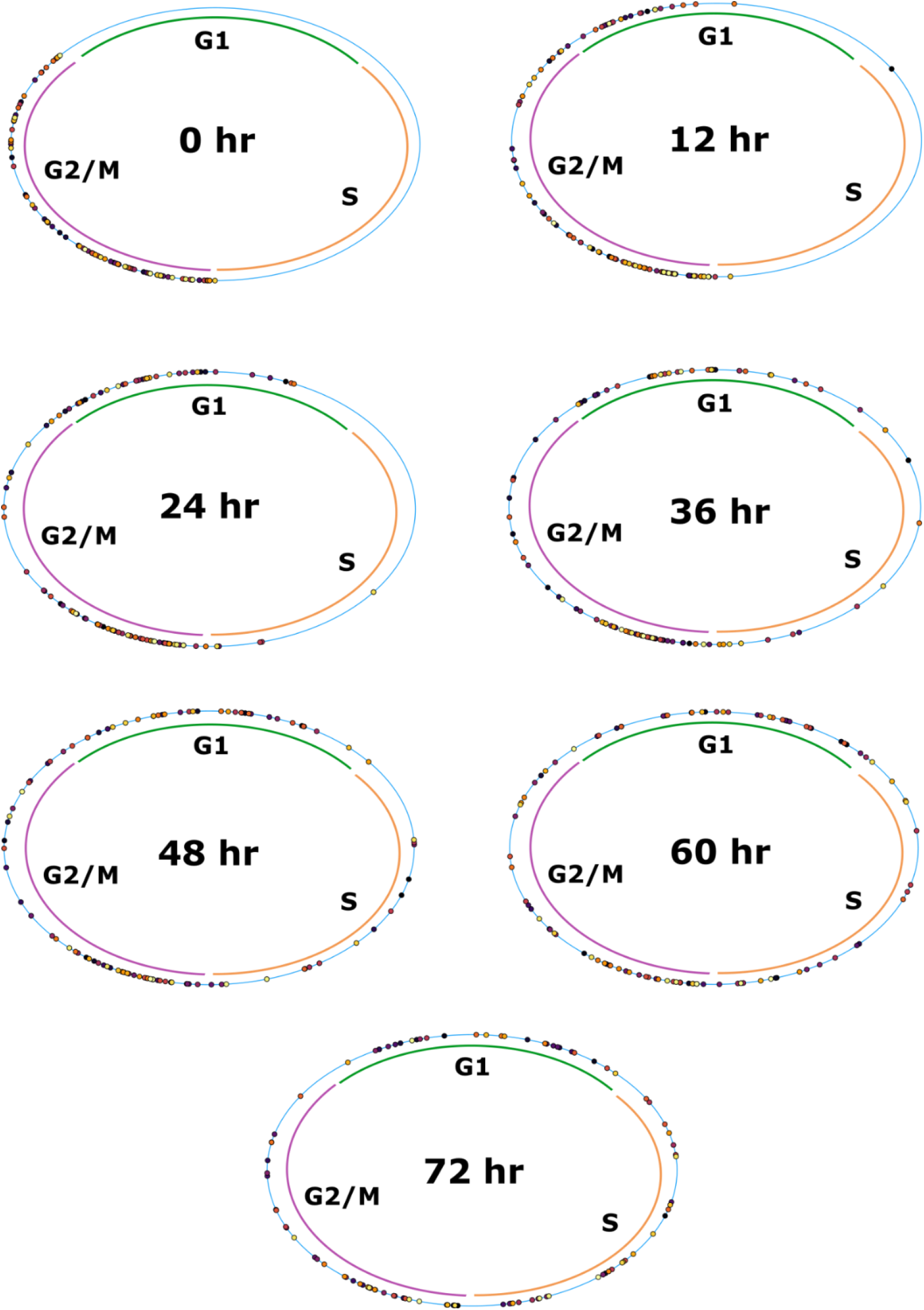
Predicted progression through the cell cycle of G2M cells The tricycle scores made based on the predictions of RNAForecaster for the 405 cells in the data are plotted on a circle diagram at each 12 hour time point.

**Supplemental Figure 8.**
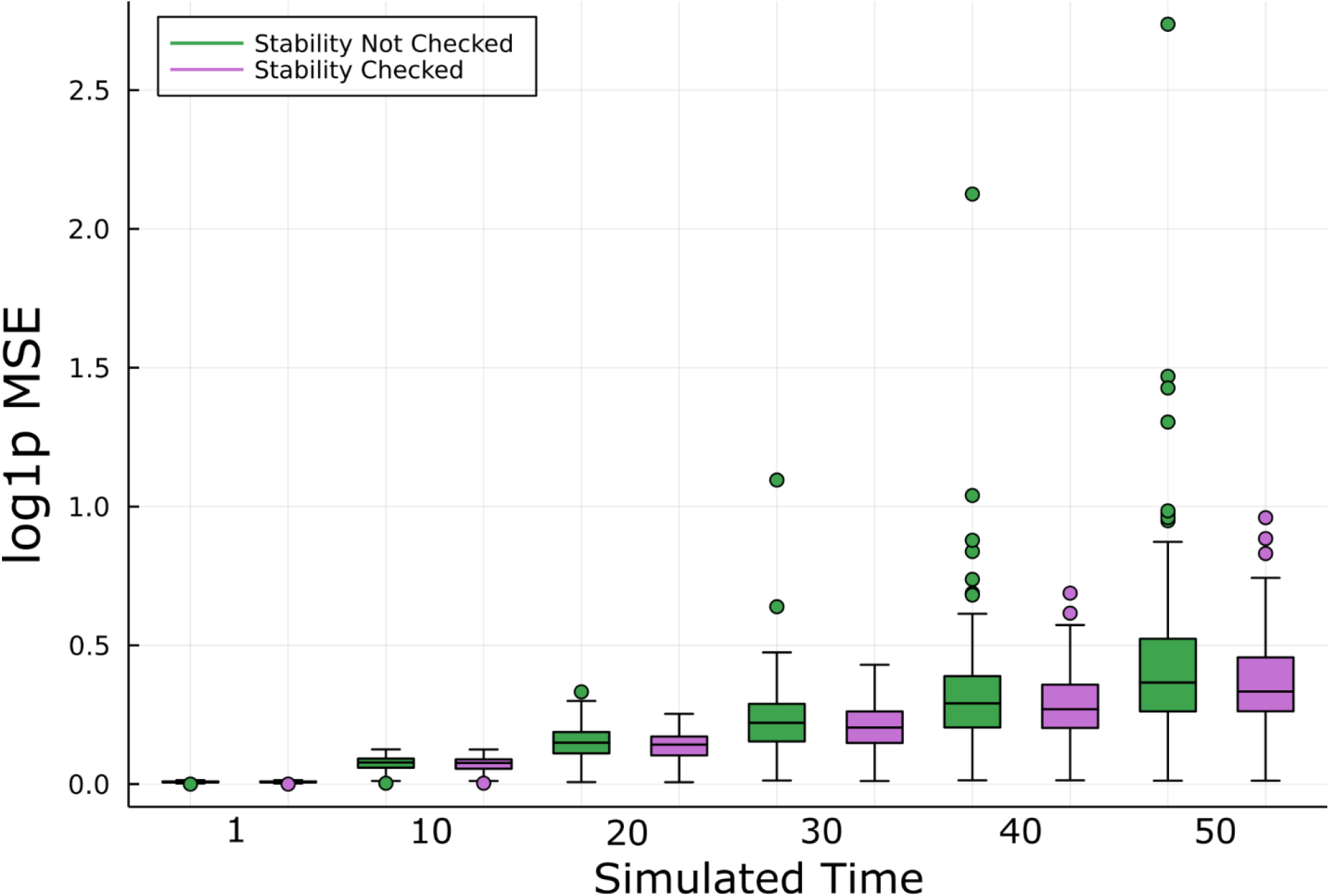
Enforcing stable predictions in simulated data Boxplots of log MSE loss on simulated data both with and without checking to see whether the model always produced predictions less than 2x the maximum expression value (the model was retrained if above the threshold). Fewer outliers, indicating potential catastrophe, are observed in the stability checked data.

